# Crops grown in mixtures show niche partitioning in spatial water uptake

**DOI:** 10.1101/2022.03.08.482442

**Authors:** Anja Schmutz, Christian Schöb

**Affiliations:** Institute of Agricultural Sciences, ETH Zurich, Universitätstrasse 2, 8092 Zurich, Switzerland; Área Biodiversidad y Conservación, Universidad Rey Juan Carlos, c/ Tulipán s/n, 28933 Móstoles, Madrid, Spain

## Abstract

1. More diverse plant communities generally produce more biomass than monocultures. This benefit of plant diversity is supposed to stem from resource partitioning of species in mixtures. Different plant species might use the resources spatially, temporally, or chemically in different ways. Along the same lines, for agricultural production crop mixtures outperform monocultures. Differences in vertical root distributions of crop species in mixtures could explain such higher yield.
2. Here we used the stable isotopes of water and a Bayesian model to investigate the spatial water uptake patterns of six different crops species and how these patterns differ depending on the crop diversity. In addition, we calculated niche overlaps of water uptake as an indicator for belowground spatial niche partitioning, compared them among the different diversity levels, and linked them to productivity.
3. The spatial water uptake pattern differed among crop species. The effect of crop diversity had a minor effect on water uptake but varied strongly depending on the crop species. Niche overlap in spatial water uptake was highest in monocultures and decreased strongly in mixtures. Furthermore, productivity in mixtures was higher compared to monoculture. Additionally, we showed that increased competition intensity leads to stronger changes in water uptake patterns.
4. *Synthesis*. We found evidence for niche partitioning of spatial water uptake, and therefore complementary spatial root distribution, and higher productivity in crop mixtures compared to monocultures. Consequently, a more efficient use of soil resources in intercropping systems might explain their yield benefits.

## Introduction

Resource partitioning occurs when different species occupy a different niche along a resource profile (or a hyper-volume in a multi-dimensional space) (Hutchinson, 1978). This means that interacting species which occupy distinct niches are more likely to coexist and have a higher fitness (Silvertown, 2004). Nevertheless, niches are not static: Recent studies demonstrated niche plasticity were species shift to other resources upon interaction with other species (Ashton et al., 2010) or depending on species diversity (Niklaus et al., 2017).

The body of evidence about the relationship between resource partitioning and species diversity effects on productivity is controversial with no/limited (Bachmann et al., 2015; Barry et al., 2020) and supporting evidence (Guderle et al., 2018; O’Keefe et al., 2019) in grassland species. Studies that investigate resource partitioning in cropping systems are lacking despite the fact that resource partitioning might be one of the explanations why mixtures (i.e. the simultaneous cultivation of two or more species in the same field) outperforms monocultures (Barry et al., 2019; Turnbull et al., 2016). Hence, simultaneously cultivated crop species that have spatially, temporally, or chemically distinct niches or have the ability to shift to an alternative resource source are likely to be more productive. In fact, field studies in grassland systems have already shown a positive relationship between resource partitioning and overyielding (i.e. higher productivity in mixtures compared to monocultures) (Mason et al., 2020; Verheyen et al., 2008).

With regard to the resource water, mixed cropping has already shown to increase water-use efficiency compared to monocropping practice (Morris and Garrity, 1993). Furthermore, mixed cropping systems show stable or even enhanced crop yield even under drought stress (Renwick et al., 2020). However, evidence about water partitioning in crop species and its relationship with species diversity are lacking. There are already some studies that investigated (spatial/temporal) water use in crop species (Asbjornsen et al., 2007; Liu et al., 2021, 2020; Ma and Song, 2018, 2016; Wang et al., 2010; Wu et al., 2018, 2020) – but they did not explore water partitioning in intercrops.

Hence, the following research questions were addressed in this study: Do different crop species use water spatially differently? Do crop species shift to spatially other water sources depending on species diversity? Do crop species grown in mixtures show spatial water partitioning? Does water partitioning in mixtures lead to higher productivity? Does competition intensity explain patterns in spatial water uptake?

A field experiment was conducted where six different crops were grown in different diversities (one, two or three species grown together). The natural abundance of stable isotopes in plant and soil water and the mixing model MixSIAR (Stock et al., 2020) was used to calculate the proportion of water from different soil layers taken up by the plants grown in different diversities. Furthermore, the niche overlap between pairs of species were calculated as an indication for niche partitioning - and was linked to productivity. Last but not least, the relationship between competition intensity and spatial water uptake was estimated to test if competition intensity can explain patterns in spatial water uptake. We hypothesised that distinct crop species differ in their spatial water uptake. Even though there is evidence about niche plasticity in response to neighbouring species and diversity in meadows and trees (Ashton et al., 2010; Niklaus et al., 2017), we would not expect any spatial plasticity of water use in crops due to the breeding (Matesanz and Milla, 2018). Hence, we assumed that niche overlaps are lower in mixtures compared to monocultures due to niche differences of the different crop species. We further expected that communities composed of species with decreased niche overlap in mixtures would result in a higher productivity. Finally, we assumed that under high (aboveground) competition plants would use more water from deeper soil layers.

## Material and methods

### Study site

The field experiment was conducted near Zurich, Switzerland (coordinates 47°26′19.917″N 8°29′58.930″E and 455 m a.s.l.). The field site is located in a temperate climate with annual mean temperature of 9.3 °C and annual mean perception of around 1000 mm (norm period 1981-2010, MeteoSchweiz). During the field experiment, temperature ranged from 0°C in the beginning of May up to 36°C at the end of June (**Fig. 1**). During June 2019, the hottest days ever were recorded, and June was the second hottest June since 1864. Furthermore, June was a dry month with only two-thirds of usual rainfall during this month (**Fig. 1**).

**Fig. 1.**
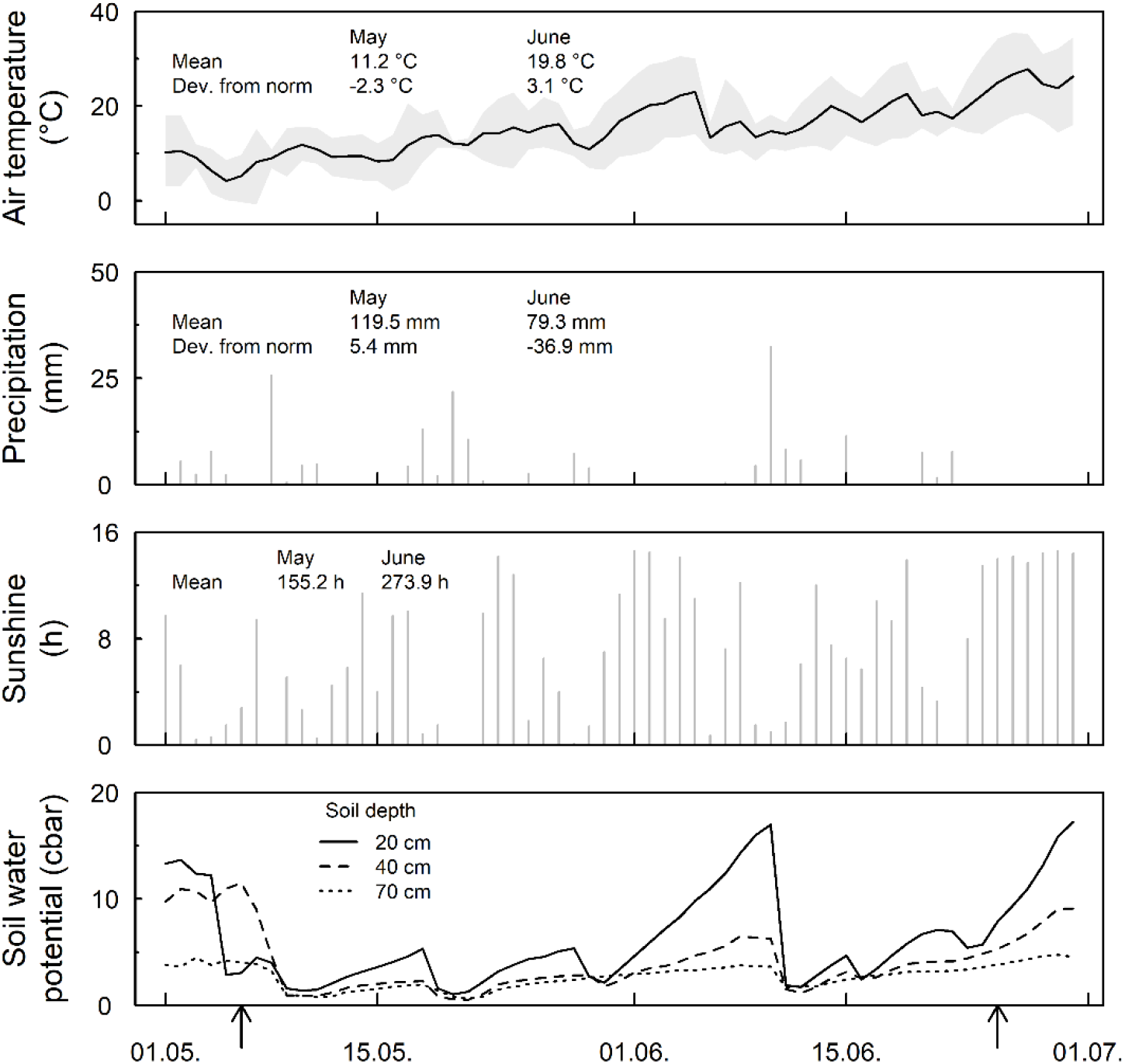
Air temperature (day mean ± day minima and maxima), precipitation, sunshine duration and soil water potential between May and June 2019. Arrows indicate sowing (6 May 2019) and the sampling event (25 June 2019). For temperature, the daily mean, minima and maxima are shown. For perception and sunshine duration, the day sum is drawn. The monthly mean of the norm period (1981-2010) and the deviation from the norm period of the months May and June 2019 are also indicated. The Data is from the MeteoSchweiz weather station Affoltern and from Kanton Zürich Bodenschutz station in Reckenholz (soil humidity) (approx. 2 km away from the field site).

The soil at the study site was a brown earth which is profound and with normal water-permeability. The soil consists of 56% sand, 18% clay, 25% silt, 3% hummus, 8% gravel and 2% stones. Soil water potential in 20 cm depth reflected precipitation, whereas in 70 cm the soil water potential remained steady (**Fig. 1**). Soil water potential can roughly be classified in the following groups: saturated-very wet (0-6 cbar), wet-moist (6-10 cbar), moist-dried (10-20 cbar), dry (20-50 cbar) and very dry (>50 cbar). During the duration of the experiment the soil in 20 cm depth was wet and only rarely drier (peak in the first part of June). This indicates that there was no water limitation during the experiment.

### Seed material

In this experiment, six crop species were used, namely spring barley (*Hordeum vulgare* var. Atrika), spring wheat (*Triticum aestivum* var. Fiorina), faba bean (*Vicia faba* var. Fanfare), pea (*Pisum sativum* var. Astronaute), linseed (*Linum usitatissimum* var. Marquise) and rapeseed (*Brassica napus* subsp. *napus* var. Campino). The seeds were purchased from a local retailer (UFA Samen) and are local varieties commonly cultivated in Switzerland. These crop species can be grouped into three functional groups: cereals including wheat and barley, legumes containing faba bean and pea and (oilseed) herbs with linseed and rapeseed.

### Experimental design

The six crop species were grown in plant communities as monocultures and mixtures with either two or three species, respectively. For the 2-species mixtures, all possible combinations among the crop species were cultivated. As for the 3-species mixtures, all possible combinations among the functional groups were grown together (**Table S1**). In total 29 different plant communities were grown. The 29 community plots together with three control plots with no plants were arranged in blocks of 4 × 8 plots. This complete set was replicated three times following a randomised-complete block design. Additionally, all crop species were grown as single plants. These single plants experienced no above- and belowground interaction during the whole cropping season. Thus, these plots were weeded regularly to prevent any growth of weeds which could interact with the crops. To exclude any above- and belowground interaction with other crops, all single plants were grown in one separate block which contained five replicates of each crop, randomly allocated within the block. All plots measured 0.5 × 0.5 m and were separated by a metal fence to a depth of approx. 30 cm.

Sowing was conducted by hand on the 6 May 2019 with the sowing density recommended by the seed retailer (210 kg ha^−1^ for wheat, 180 kg ha^−1^ for barley, 250 kg ha^−1^ for faba bean, 275 kg ha^−1^ for pea, 60 kg ha^−1^ for linseed and 6 kg ha^−1^ for rapeseed). The sowing ratio (%) was 50:50 and 33:33:33 for 2-species and 3-species mixtures, respectively, except for rapeseed which had 20% in both 2-species and 3-species mixtures (**Table S1**). Effective plant counts were conducted on the 5 June 2019 in all plots. Since the distinguishment between barley and wheat in the 2-species mixture was difficult at this early growth stage, the total plant count in these plots was divided by two as the sowing ratio was 50:50.

### Water sampling

Sampling for natural ^18^O and ^2^H abundance took place on 25 June 2019. In four blocks (three with the community plots and one with single plants), two control plots per block were randomly selected for soil core sampling. Soil cores were taken with a core sampler and the soil was collected in the following depths: 0, 5, 10, 15, 20, 30, 50 and 75 cm. These empty plots were sampled as control to reduce any effect of crops on soil water (i.e. hydraulic lift). Furthermore, the small scale of the plots would also not exclude any possible diffusion of soil water between plots. Hence, due to these motives and feasibility constraints to sample soil cores in all the plots, empty plots were chosen in each block for soil water sampling and not the corresponding plots with plants.

For the examination of source water, water extracted from the root crown has already shown to match the source water best (Barnard et al., 2006). Hence, for the community plots the root crown of one random individual per species and plot was collected. For linseed, three individuals per plot were pooled, because one individual did not contain enough water for extraction. For the single plants, three plants per crop were randomly chosen for sampling. The soil and plant samples were collected in 12 ml glass vials, closed with a chlorobutyl rubber septum cap (Exetainer©, Labco Limited) and stored at −20°C until further processing.

### Isotope analysis

Water from samples was extracted following the Cryogenic vacuum method (Dalton, 1988). Subsequently, extracted water was analysed for hydrogen and oxygen isotopic composition (δ^2^H and δ^18^O) following the protocol by Werner and Brand (2001). The results were normalised to VSMOW (Coplen, 1988) and are expressed in the δ-notation (**Eq. 1**).

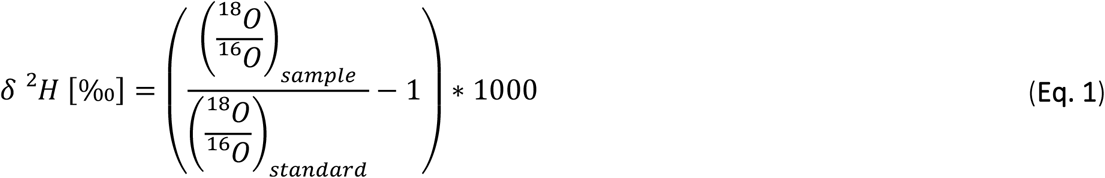

### Biomass sampling

During the water sampling, the shoot from the sampled plants were collected in paper bags and dried at 80°C for at least 48h. Subsequently, the dry biomass was weighed.

### Data analysis

The complete analysis was carried out in R version 4.0.2 (R Core Team, 2021).

For the relations between δ^18^O and δ^2^H in soil water and the δ^18^O and δ^2^H along the soil profile linear mixed models were conducted. In the first model, δ^2^H was the response variable, δ^18^O the predictor and the plot the random term. For the δ^18^O and δ^2^H along the soil profile, soil depth was the response, δ^18^O and δ^2^H (second-order polynomial regressions), respectively, were the predictors and plot the random term.

To quantify the proportion of water uptake (PWU) of the different plants grown in the different diversities, the Bayesian isotope mixing model MixSIAR (Stock et al., 2018) was used. This model allows to calculate the proportion of sources (here the soil water from the specific soil depth) of unknown mixtures (here the sampled plant xylem water). In advance, the measured δ^18^O and δ^2^H values from the specific soil depths were combined into the following soil layers: shallow (0-10 cm), middle (15-30 cm) and deep (>50 cm). This spatial stratification of the source allows a better model inference and analysis (Phillips et al., 2005; Stock et al., 2018). Differences between the soil layers were tested with student’s t-tests that compare the three soil layers in all possible combinations in both δ^18^O and δ^2^H values. With MixSIAR, two models were computed. Model 1 describes the PWU from the six crops in the different diversity levels (fixed factor) with the replicates as random factor (**Fig. 3**). Model 2 was run for each replicate (specific species composition as fixed factor, no random factor). This model describes how the water uptake differs when specific species are grown together (**Fig. S1**). Each model was run with a chain length of 300’000, burn-in of 200’000, thinning of 100 and with three chains (“long” run in MixSIAR).

Furthermore, niche overlap in water uptake was calculated from the overlap of the posterior distribution estimated in model 1 (**Fig. S2**). For this, the overall of the kernel density estimates between all the possible hypothetical crop species pairs of all the different diversities and soil layers was calculated with the R package *overlapping* version 1.6 (Pastore, 2018). The use of overlapping estimates of two or more kernel densities has been suggested for calculating niche overlap rather than only compare means or use the geometrical overlap of kernel densities (Pastore, 2018; Pastore and Calcagnì, 2019; Swanson et al., 2015). Subsequently, a hierarchical cluster analysis (*hclust* function in R) was applied to the overlapping values. This analysis reveals patterns in niche overlaps in the different diversity levels, soil layers and in the hypothetical species pairs. Furthermore, the sum of niche overlaps across all three soil layers was estimated and paired t-tests between species diversities were computed with the hypothetical species pairs as grouping variable. This analysis was only done for the species grown in community since single plant niche overlaps were far from normally distributed - and for comparison with productivity, the community plots were more relevant.

Productivity (g/m^2^) of species grown in community (monoculture, 2- and 3-species mixture) was calculated from the biomass collected during sampling and the number of plants counted after germination and the sowing ratio (productivity = biomass * count / sowing ratio * 4). Subsequently, the mean productivity for each species in each diversity level was calculated and the productivity of crop pairs was estimated by the sum of all possible combinations of hypothetical species pairs in the separate diversity levels. Paired t-tests between the species diversities were computed to estimate differences of productivity of hypothetical species pairs (same as for the niche overlaps). In addition, a linear mixed model (LMM) was computed with the productivity as response, species diversity and niche overlap as predictors and the hypothetical species pairs as random term. This LMM gives information about the relationship between productivity and niche overlap of hypothetical species compositions at different diversity levels.

Furthermore, as a measure of the intensity of plant-plant interactions, the relative interaction index (RII) with the biomass of the plant when grown in community (B_c_; monoculture, 2- or 3-species mixture, respectively) and when grown without any other plant (B_s_; single plants) was calculated (Armas et al., 2004) (**Eq. 2**).

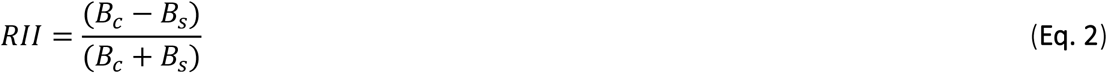

The RII gives information about how the biomass changes upon interaction with neighbours, RII<0 indicates competition (negative interaction) whereas RII>0 indicates facilitation (positive interaction). RII was previously transformed to allow for square-root transformation (RII_t_ = (RII + 1) / 2). Subsequently, a LMM was computed to test if RII for the different crop species differed depending on the species diversity. The square-root transformed RII_t_ for each species separately was the response variable, the block, diversity (monoculture vs mixture) and the mixture diversity (2- vs 3-species mixture) were the explanatory variables and the species composition was set as random term.

For the relationship between RII and mean PWU (estimated in model 2), a generalised linear mixed model (GLMM) with beta distribution was computed. Prior, mean RII of the three replicates were calculated. Since the mean PWU of the three soil layers are not independent, a separate model for each soil layer was implemented. The mean PWU was the response variable and the RII, species diversity (monoculture, 2- and 3-species mixture) and their interaction were the explanatory variables, and the species composition was set as random term.

## Results

### Isotopic signature of soil water

The δ^18^O and δ^2^H values from water extracted in the specific soil depths were correlated (P < 0.001) and showed δ^18^O values between −1‰ and −11‰ and δ^2^H values between −27‰ and - 77‰ (**Fig. 2a**). When looking at the isotopic signature along the soil profile the following pattern is visible: The topsoil is isotopically enriched, and the isotopic signatures of δ^18^O and δ^2^H decrease with increasing soil depth (**Fig. 2b-c, Table S2**).

**Fig. 2.**
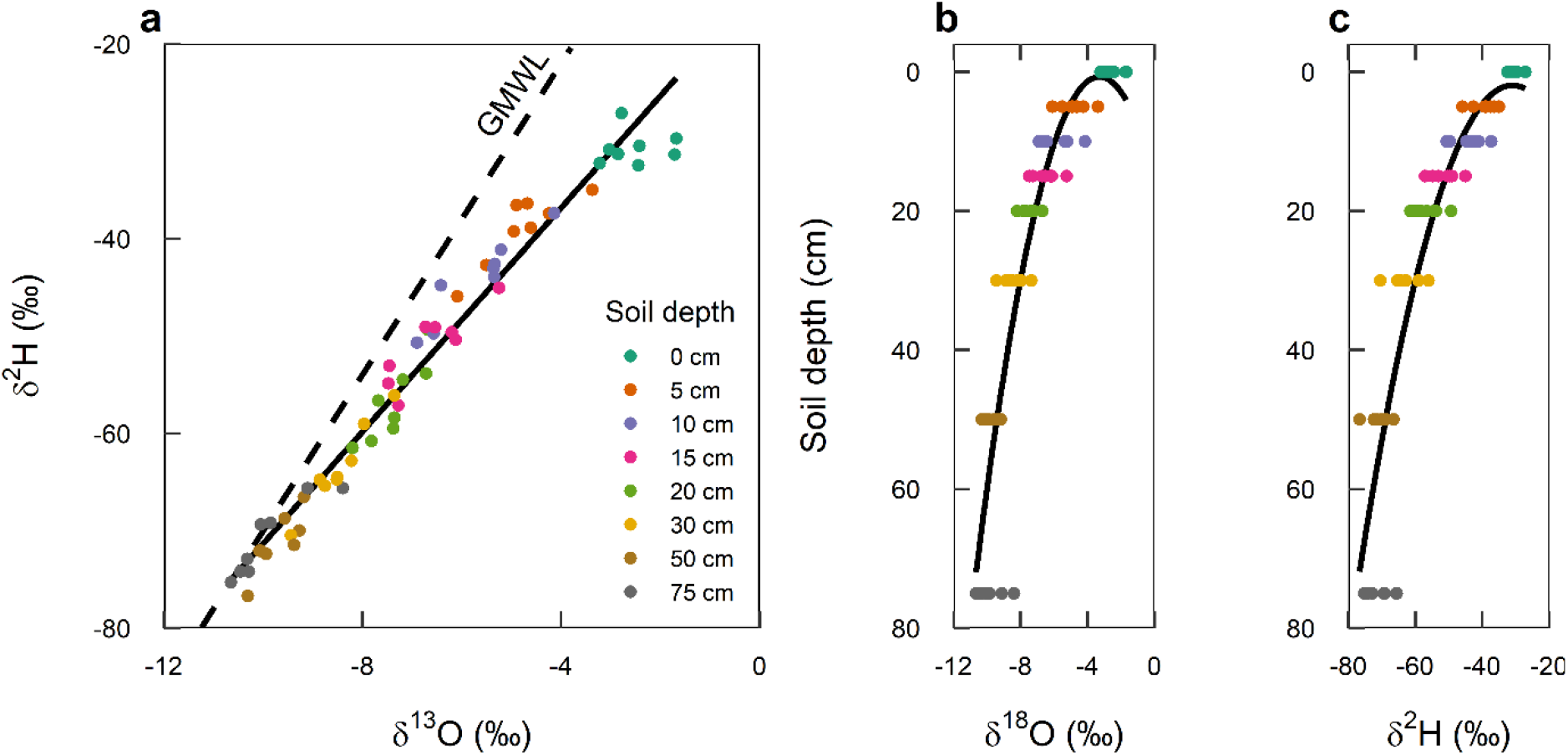
Isotopic signature of δ^18^O and δ^2^H (‰) from water extracted at specific soil depths. (**a**) Relationship between δ^18^O and δ^2^H. Water extracted from the soil depth are coloured accordingly (points in different colours). The regression line of the soil water is shown (solid line; y = −13.8 + 5.7x; P < 0.001; N=64). The global meteoric water line is also shown (GMWL; dashed line; y = 10 + 8x). (**b**) δ^2^H values along the soil profile. Shown is the regression line (y = −26 + 165 - 48x^2^; P < 0.001) and the individual samples per soil depth (coloured points; N=8). (**c**) δ^18^O values long the soil profile. Shown is the regression line (y = −26 + 162x - 65x^2^; P < 0.001) and the individual samples per soil depth (coloured points; N=8).

**Fig. 3.**
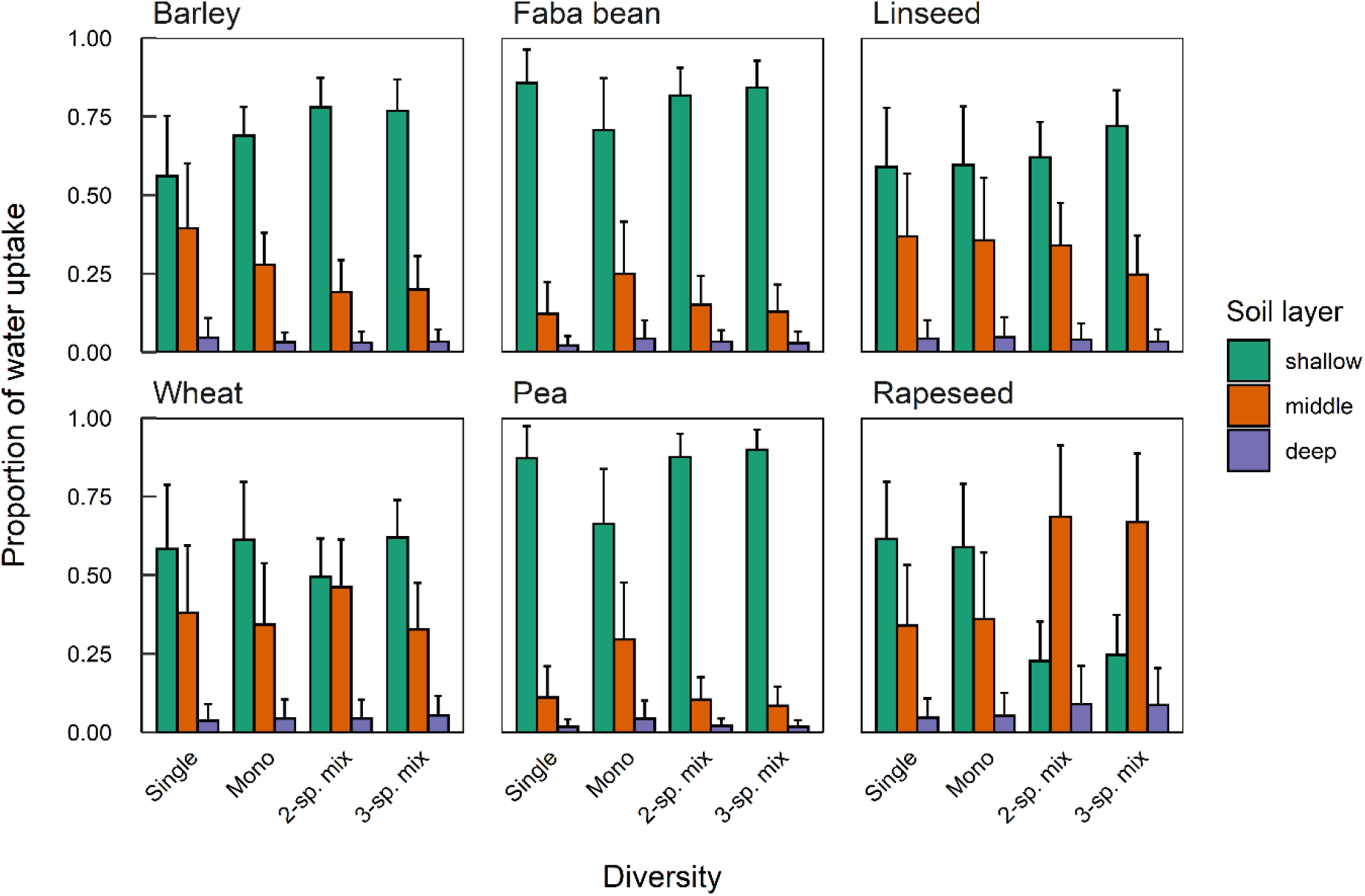
Proportion of water uptake from the different soil layers by the six crops barley, wheat, faba bean, pea, linseed, and rapeseed grown as single plant, in monoculture and in 2- and 3-species mixtures (mean ± standard deviation of the mean). The proportions were calculated with the mixing model MixSIAR (model 1; species*diversity as fixed factor, species composition as random factor).

The different soil layers (shallow, middle and deep) showed significantly different isotopic signatures in both δ^18^O and δ^2^H (P < 0.001 between all soil layers, data not shown).

### Proportion of water uptake in the different soil layers

In our experiment, most of the water was taken up in the shallow soil layer (**Fig. 3**). The importance of the middle soil layer (15-30 cm) varies mostly between the species but also within the species between diversities (**Fig. 3**). In general, the deep soil layer (>50 cm) was not an important water source. For the cereals, barley and wheat, the shallow and middle soil layers were both important sources of water. For wheat, the proportion remained unchanged across all diversities whereas in barley a trend toward a higher PWU with increasing species diversity is visible. For both legumes, faba bean and pea, most of the water was taken up in the shallow soil layer and did not change among the diversity levels. For linseed, shallow and middle soil layer were both an important source of water. On the other hand, rapeseed showed a clear shift from shallow to middle soil layer, when grown in mixtures compared to when grown as single plant or in monoculture.

### Niche overlap in spatial water uptake

The cluster analysis revealed three distinct clusters of PWU among the different diversity levels and soil layers (**Fig. 4**, x-axis): the first cluster with the monocultures independent of soil layer (D1) and the deep soil layer independent of the diversity level (L3), the second cluster with the shallow and middle soil layer (L1 and L2) in single plants (Ds) and the third cluster with the shallow and middle soil layer (L1 and L2) in 2- and 3-species mixtures (D2 and D3). The first group including all monocultures was characterised with high overlap values (≥ 0.5). In the second cluster including the single plants, overlap values depended strongly on the hypothetical species combination. When the two legumes (faba bean and pea) would be combined with the other species, overlap values would be low (<0.3) whereas these values would be higher among the other species (>0.8). The third cluster including all the mixtures shows low to very low overlap values (<0.5, with some exceptions). This indicates that niche overlap is lowest in the mixtures.

**Fig. 4.**
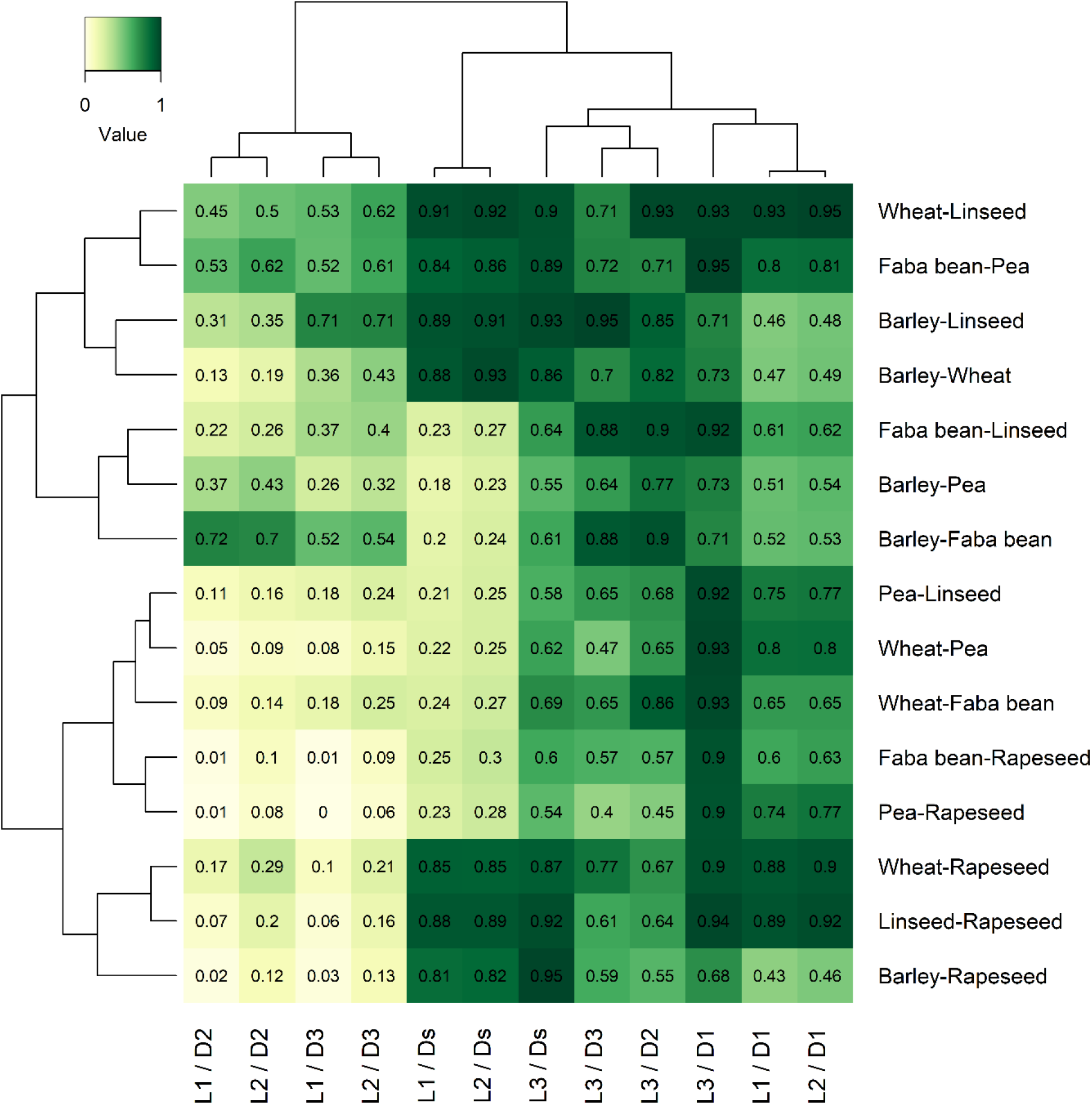
Heatmap of the estimated overlaps from the posterior densities estimated by model 1. Overlaps were calculated between all the possible combinations of species and for the interactions of soil layer (L; 1=shallow, 2=middle, 3=deep) and the diversity (D; s=single plant, 1=monoculture, 2=2-species mixture, 3=3-species mixture). Overlap values range from zero to one and are also indicated in the corresponding squares. Histograms show hierarchical clustering (*hclust* function in R).

The clustering of the species combinations revealed two clear clusters (**Fig. 4**, y-axis). These two clusters mainly differed in their overlap value estimated for the two mixtures (D2 and D3) in the two upper soil layers (L1 and L2). The cluster with higher overlap values included all mixtures with barley or faba bean, except when they are combined with rapeseed and faba bean-wheat, and the combination wheat-linseed. Overlap values of rapeseed with any other species were low (<0.29), as are the combinations of wheat with faba bean and pea, and pea-linseed. To summarise, niche overlap in water uptake is lowest in the mixtures – especially when the combination includes rapeseed or wheat with legume.

### Linking niche overlap to productivity

Productivity of hypothetical species pairs was significantly higher in the 2- and 3-species mixtures compared to the monoculture (**Fig. 5a**). Analogous, the sum of niche overlaps across all soil layers was significantly higher in monocultures when compared to the two mixtures (**Fig. 5b**). Furthermore, there was no relationship between niche overlap and productivity in any species diversity (**Fig. S3**).

**Fig. 5.**
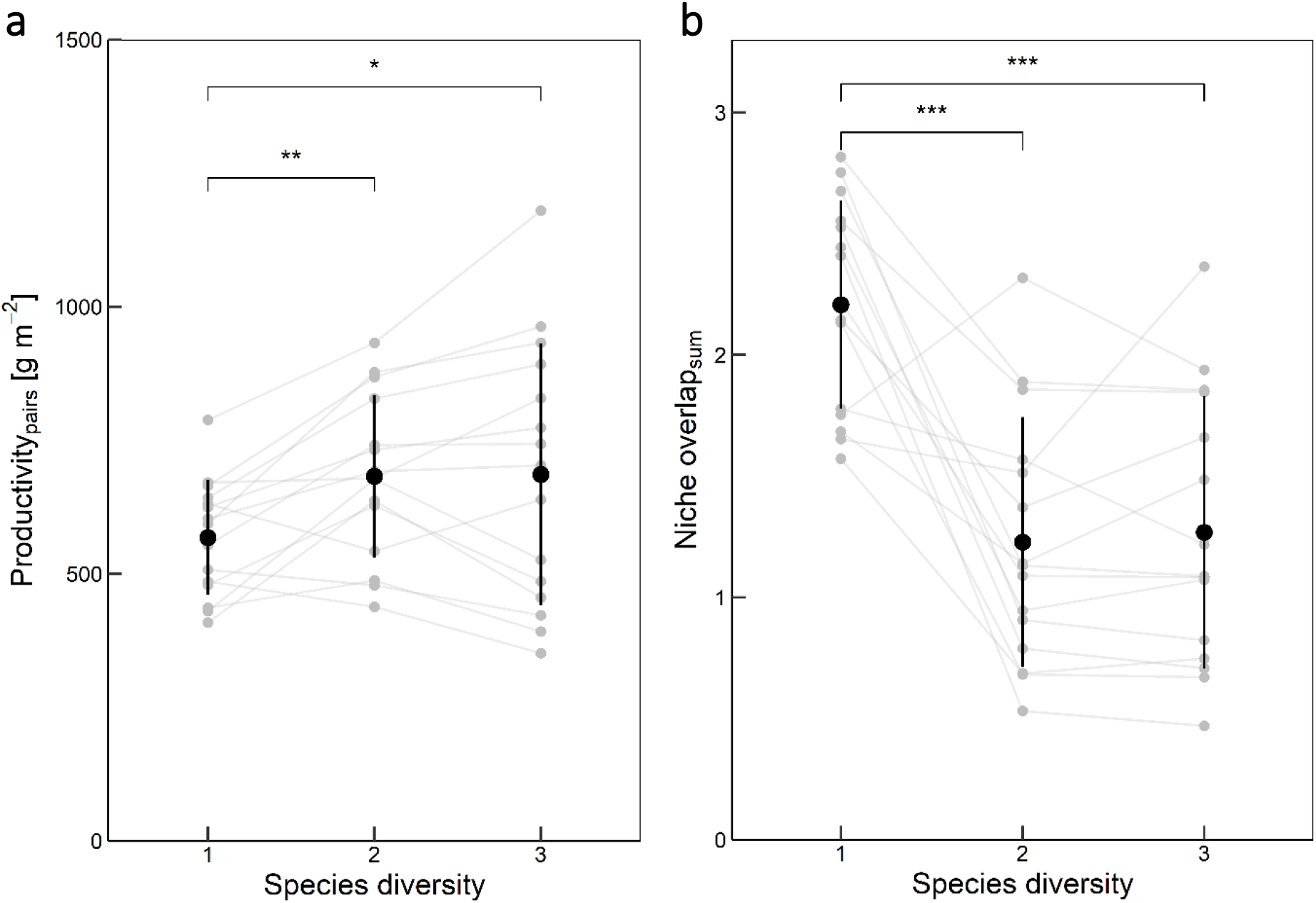
Productivity (a) and sum of niche overlap across all soil layers (b) of hypothetical species pairs grown in different species diversities. Shown are the single data points (grey points), the connection between paired data points (hypothetical species pairs; grey lines) and the mean ± standard deviation of the mean (black point and line). Significant differences of a paired t-test are indicated with asterisk and brackets. Significance code corresponds to *: P<0.05, **: P<=0.01, ***: P<=0.001.

### Relationship between competition intensity and spatial water uptake

RII differed among species (**Fig. 6**). Rapeseed showed the highest reduction of biomass upon interaction with neighbours (approx. −0.85), closely followed by barley (approx. −0.7), wheat and linseed (both approx. −0.6). Competition was relatively small for the two legumes faba bean and pea. Nonetheless, the RII, as it was the case for all species, was significantly smaller than zero (p<0.001, **Table S3**). Regarding species diversity, only linseed showed a response to increasing crop diversity with lower RII (higher competition) in the two mixtures compared to the monoculture (p=0.047, **Table S3**).

**Fig. 6.**
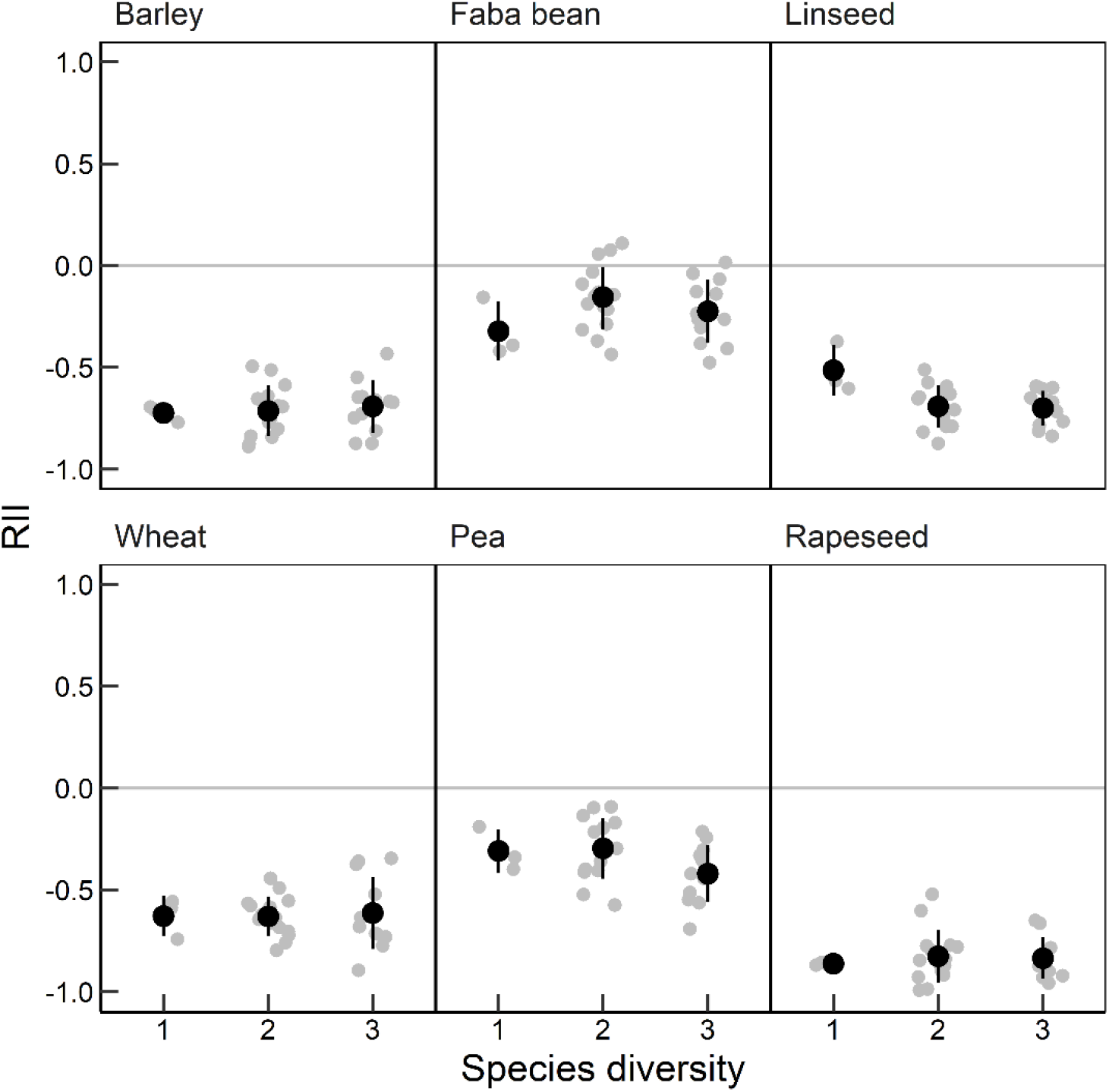
Relative interaction index (RII) calculated as the relative change in biomass between individuals grown in community (monocultures and 2- and 3-species mixtures) and as single plant without neighbours. Drawn is the mean ± the standard deviation of the mean (black points and lines) and the individual data points (grey points).

To investigate if patterns in water uptake change upon interaction intensity, the relationship between mean PWU and RII was estimated. The GLMM and subsequent type-III ANOVA indicated significant effects of RII on mean PWU in the shallow and middle soil layer (**Fig. 7, Table S4**). The interaction between RII and species diversity was not significant (**Table S4**). In other words, more intense competition led to a lower proportion of water uptake in the shallow soil layer, whereas in the middle soil layer, more intense competition led to a higher proportion of water uptake. This indicates that upon more intense competition, crop species use more water from deeper soil. Nevertheless, the relationship between mean PWU and RII is mainly driven by differences of RII between species since the effect of diversity on RII within the species is small (only one significant effect of diversity on RII in linseed; **Table S3**).

**Fig. 7.**
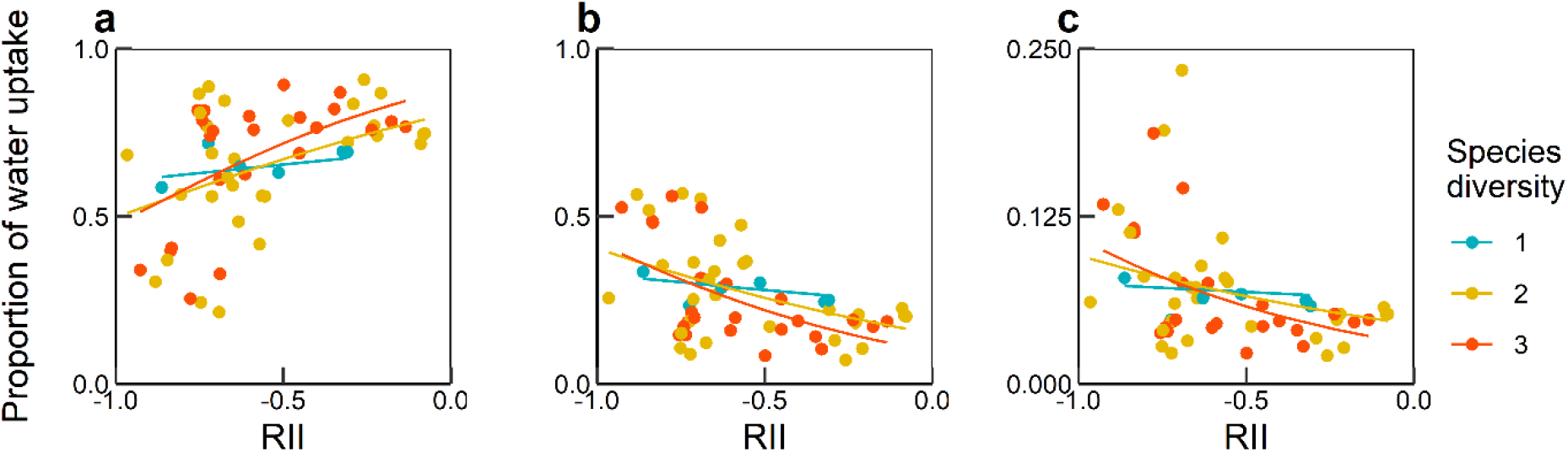
Relationship between mean proportion of water uptake (from model 2) and relative interaction index (RII) in the (a) shallow, (b) middle and (c) deep soil layer. shown are the single data points and the prediction of the generalised linear mixed model in the different species diversities (1-, 2- and 3-species). Note the different scale of the y-axis in (c).

## Discussion

The methods applied, which included stable isotopes of water and subsequent application of mixing models, allowed to investigate the spatial water uptake of six different crop species grown in monocultures and mixtures and to assess water partitioning between species pairs.

### Isotopic signature along the soil profile and soil water potential

The soil profile in this study showed clear enrichment in heavy isotopes (i.e. higher δ^18^O and δ^2^H) in the top soil and continuous decrease with soil depth (**Fig. 2**). Furthermore, when comparing the global meteoric water line, the regression line from collected stable isotopes in this study are less enriched in δ^2^H compared to δ^18^O. This effect is due to evaporation (Cappa et al., 2003) in the top soil. Temperature, precipitation, and soil water potential data support these findings (**Fig. 1**). Since the last rain event in the first part of June, topsoil (20 cm) was continuously drying out – but only to a point where the soil can be classified as moist at the time of data sampling.

At that stage it is important to note that no soil moisture data is available for the study site directly or the different plots which could explain patterns in spatial water uptake. However, the plots are arranged on a small scale and thus diffusion between plots cannot be excluded. Hence, large differences in soil moisture distribution between diversities and/or species compositions are not expected. Therefore, we assumed that observed patterns in spatial water uptake are rather a product of the diversity and/or species composition and not of the soil moisture distribution in the soil.

### General patterns in spatial water uptake and the relationship to root distribution

The shallow (0-10 cm) and middle (15-30 cm) soil layers were the most relevant water sources for the inspected crop species (**Fig. 3**). These findings are in accordance to the plant water uptake globally in the temperate climate zone (Amin et al., 2019) and grassland species that were grown in different diversities (Guderle et al., 2018). Nevertheless, spatial water uptake depends strongly on plant growth stage (Ma and Song, 2018, 2016; Wang et al., 2017; Wu et al., 2018). These studies show that crop species used a big proportion from shallow soil (0-20 cm) in early stages and shift to deeper soil layers just before flowering. In our study, the crops were in the stage of heading (wheat and barley), flowering (faba bean), and early fruit development (pea, linseed and rapeseed). During these stages, the roots are expected to be fully developed (Weaver, 1926). Plant roots therefore explore the greatest possible soil area and depth.

In fact, it is suggested that water uptake and root distribution are tightly coupled (Gardner, 1964). However, results from field studies are controversial with evidences of both positive and negative relationships between PWU and root length density (Guderle et al., 2018; Liu et al., 2021; Ma and Song, 2018). The six different crop species indeed have distinct rooting patterns (Fan et al., 2016; Kutschera et al., 2009). For instance, rapeseed shows a higher root length density compared to other oilseed and pulse crops (among others linseed and pea) (Liu et al., 2011), whereas faba bean has in general a shallow root system (Li et al., 2006). These distinct rooting patterns are reflected in the observed differences in spatial water uptake of the crop species when grown as single plants. Furthermore, due to the gradient of water isotopes along the soil profile (**Fig. 2**), water isotopes can be used to indirectly explain root distribution especially when comparing between different species diversities.

### The effect of crop diversity on resource partitioning and plasticity in water uptake

Literature suggests that resource partitioning is stronger in more diverse communities due to differences in resource uptake between species (Barry et al., 2019). For example, when species are grown in monoculture, the whole community uses the same resources. Nevertheless, when two or more species are grown together, species differ in resource uptake and, hence, available resource uptake by the community can be exploited more efficiently. On the other hand, results from field experiments suggest plasticity in spatial water uptake in response to species diversity in grassland communities (Guderle et al., 2018). This might be due to plant root plasticity in response to neighbouring plants (Callaway et al., 2003) and spatial plant roots segregation to avoid competition (Schenk et al., 1999). With regard to crop species, domestication of crops might have reduced the plasticity, for example in water use (Matesanz and Milla, 2018) – even though plants are suggested to be highly plastic in general (Sultan, 1987).

Results in this study suggest plasticity of water uptake in response to species diversity (**Fig. 4**). Niche overlaps of hypothetical species pairs were reduced in the mixtures compared to monocultures (**Fig. 5**). This indicates shifts to different water sources when grown with other species. Particularly, hypothetical species pairs with wheat or rapeseed in mixtures showed the strongest reduction in niche overlaps. Nonetheless, plasticity does not only decrease niche overlaps (Lipowsky et al., 2015). Nonetheless, in our system only a few hypothetical species pairs showed increased niche overlaps (e.g. barley-faba bean in 2-species mixture and barely-linseed in 3-species mixture).

Additionally, literature suggests that both selection and plasticity are important for niche partitioning (Meilhac et al., 2020). Since seeds in this study originated from a commercial seed supplier, crop species did not experience prior selection to e.g. mixed cropping. Thus, observed differences between monocultures and mixtures were mainly driven by plasticity (**Fig. 5**).

For crop production, plasticity might be relevant (Nicotra and Davidson, 2010) especially for resource partitioning in mixtures (Zhu et al., 2015). It is proposed that future breeding programs should emphasise on adaptive plasticity of crop species (Brooker et al., 2022; Milla et al., 2017) such as root plasticity (Schneider and Lynch, 2020) especially in more variable environments and due to climate change (Matesanz et al., 2010; Nicotra et al., 2010).

### Relationship between resource partitioning and productivity

Increased resource portioning in mixtures can explain why mixtures perform better than monocultures (Mason et al., 2020; Verheyen et al., 2008). In this study, we found lower niche overlaps and higher productivity in mixtures compared to monocultures (**Fig. 5**). Nevertheless, no relationship was present between niche overlaps and productivity in the different species diversities (**Fig. S3**). These results suggest that spatial water partitioning was not the (main) driver that led to higher productivity in mixtures. Temporal, spatial or chemical partitioning of other resources such as nitrogen (Ashton et al., 2010; Engbersen et al., 2021) or light (Mason et al., 2020) could also explain higher productivity in mixtures. In contrast, another study suggests that belowground resource uptake might not explain the positive relationship between species diversity and productivity (Jesch et al., 2018). Another explanation could be that the measured spatial water uptake at a specific time point cannot explain higher productivity. This is also a limitation of this study: Looking at only one time point might not be enough to study the impact of resource uptake on productivity (Trinder et al., 2013). A method which integrates both spatial and temporal water uptake might be more suitable to study the relevance of resource complementarity (Jesch et al., 2018).

### Competition intensity and spatial water uptake

In our system, competition (i.e. RII) was very strong (**Fig. 6**). All species showed a reduction of biomass when grown in communities. However, literature suggests that more diverse communities show reduced competition due to niche partitioning (Loreau and Hector, 2001). In our system, the effect of crop diversity on competition intensity was limited, and even showed the opposite direction (stronger competition with increasing crop diversity as seen in linseed). (Aboveground) competition and spatial water uptake also showed a significant relationship: upon more intense (aboveground) competition, the crop species used less water from shallow soil and more from deeper soil (**Fig. 7**). Hence, plant roots seem to escape into deeper soil layers when aboveground competition is strong. Since the effect of species diversity on RII was limited (**Table S3**), the relationship between (aboveground) competition and spatial water uptake is mainly driven by difference of RII between species and the water uptake by the crop species grown in the different diversities.

### Conclusion and further research

In this study we demonstrated that the investigated crop species used water spatially differently. We further showed that crop species have plasticity in spatial water uptake when grown with other species (i.e. mixtures). Moreover, water partitioning and productivity were greatly increased in mixtures compared to monocultures. Nonetheless, water partitioning does not seem to (fully) explain the higher productivity in mixtures. Furthermore, with higher (aboveground) competition crop species used more water from deeper soil layers.

It would now be interesting to investigate different irrigation and/or drought scenarios to test water partitioning in the different diversities and with different water availabilities. This might shed more light on drought resilience of crop mixtures – which would be relevant especially in rainfed agricultural areas and where drought events might become more frequent due to climate change (IPCC, 2014).

## Author contributions

AS planned and conducted the experiment, analysed the data, and wrote the manuscript. CS gave inputs to the experimental design, data analysis and the manuscript.

## Data Availability Statement

The data will be published via an open-access repository zenodo or similar.

## Acknowledgments

We would like to thank Anna Bugmann and Jianguo Chen for their help in the field and Agroscope for the use of their field. Thank you also to Michael Thieme for the constructive brainstorming sessions. The study was part of the DIVERSify project funded by the European Union’s Horizon 2020 research and innovation programme under grant agreement No. 727284. We also thank the editor and the two anonymous reviewers for the very constructive comments on a previous version of this manuscript.

