## Supplementary 1 for "Crops grown in mixtures show niche partitioning in spatial water uptake"

**Supporting Information**. Schmutz A. and C. Schöb. 2022. Crops grown in mixtures show niche partitioning in spatial water uptake.

**Table S1**. Overview of the different diversity levels (community, diversity, and mixture diversity), the species combinations and the sowing ratios.

| **community** | **diversity** | **mixture diversity** | **species composition** | **sowing ratio** |
| --- | --- | --- | --- | --- |
| single plants |  |  | wheat | 1 individual |
|  |  |  | barley | 1 individual |
|  |  |  | faba bean | 1 individual |
|  |  |  | pea | 1 individual |
|  |  |  | linseed | 1 individual |
|  |  |  | rapeseed | 1 individual |
| community | monoculture |  | wheat | 1 |
|  |  |  | barley | 1 |
|  |  |  | faba bean | 1 |
|  |  |  | pea | 1 |
|  |  |  | linseed | 1 |
|  |  |  | rapeseed | 1 |
|  | mixture | 2-species mixture | wheat-barley | 0.5:0.5 |
|  |  |  | wheat-faba bean | 0.5:0.5 |
|  |  |  | wheat-pea | 0.5:0.5 |
|  |  |  | wheat-linseed | 0.5:0.5 |
|  |  |  | wheat-rapeseed | 0.8:0.2 |
|  |  |  | barley-faba bean | 0.5:0.5 |
|  |  |  | barley-pea | 0.5:0.5 |
|  |  |  | barley-linseed | 0.5:0.5 |
|  |  |  | barley-rapeseed | 0.8:0.2 |
|  |  |  | faba bean-pea | 0.5:0.5 |
|  |  |  | faba bean-linseed | 0.5:0.5 |
|  |  |  | faba bean-rapeseed | 0.8:0.2 |
|  |  |  | pea-linseed | 0.5:0.5 |
|  |  |  | pea-rapeseed | 0.5:0.5 |
|  |  |  | linseed-rapeseed | 0.8:0.2 |
|  |  | 3-species mixture | wheat-faba bean-linseed | 0.33:0.33:0.33 |
|  |  |  | wheat-faba bean-rapeseed | 0.4:0.4:0.2 |
|  |  |  | wheat-pea-linseed | 0.33:0.33:0.33 |
|  |  |  | wheat-pea-rapeseed | 0.4:0.4:0.2 |
|  |  |  | barley-faba bean-linseed | 0.33:0.33:0.33 |
|  |  |  | barley-faba bean-rapeseed | 0.4:0.4:0.2 |
|  |  |  | barley-pea-linseed | 0.33:0.33:0.33 |
|  |  |  | barley-pea-rapeseed | 0.4:0.4:0.2 |

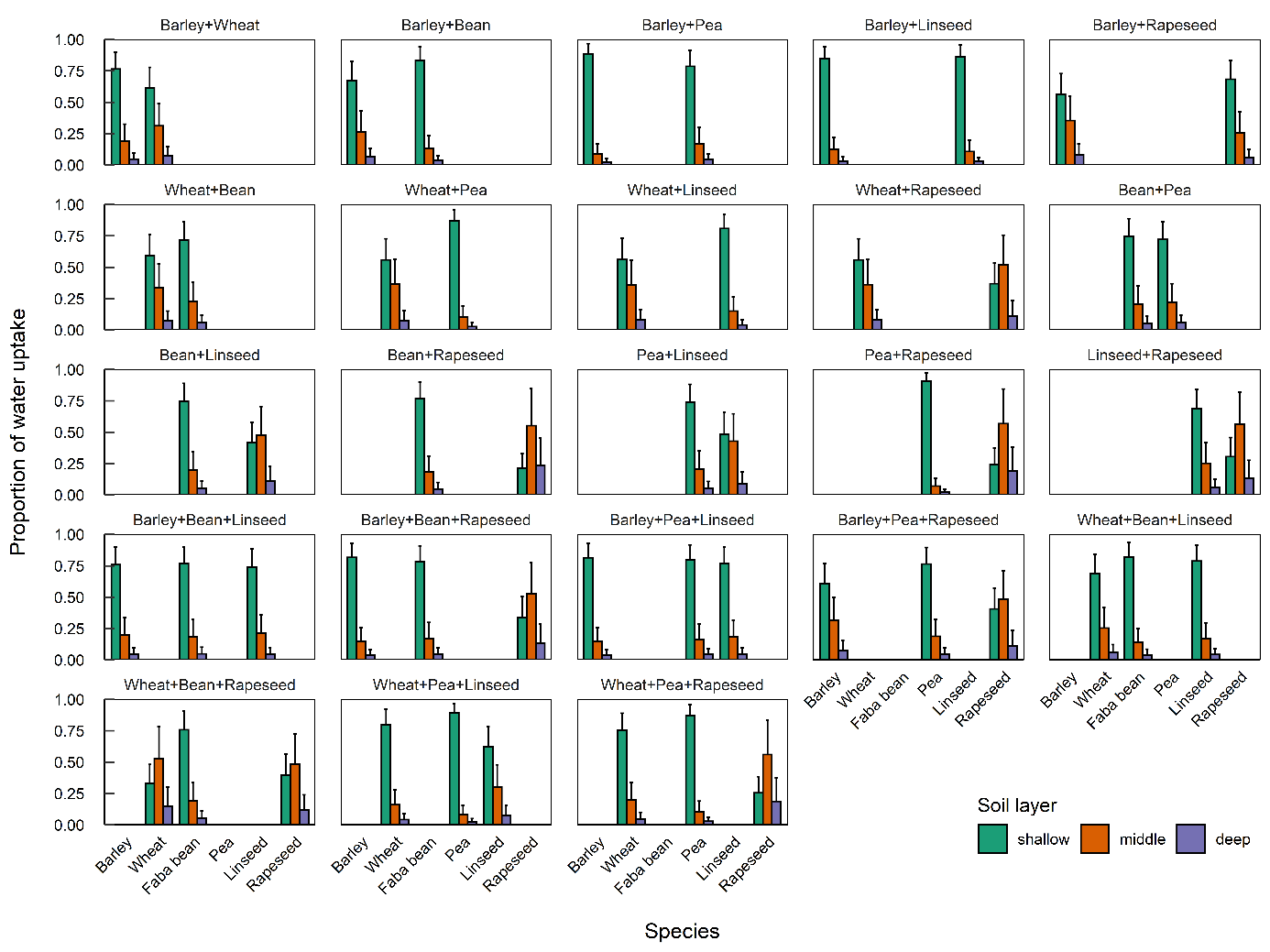

**Fig. S1**. Proportion of water uptake from the different soil layers by the six crops barley, wheat, faba bean, pea, linseed and rapeseed grown in 2- and 3-species mixtures (mean ± standard deviation of the mean). The proportions were calculated with the mixing model MixSIAR (species*diversity*species composition as fixed factor, no random factor).

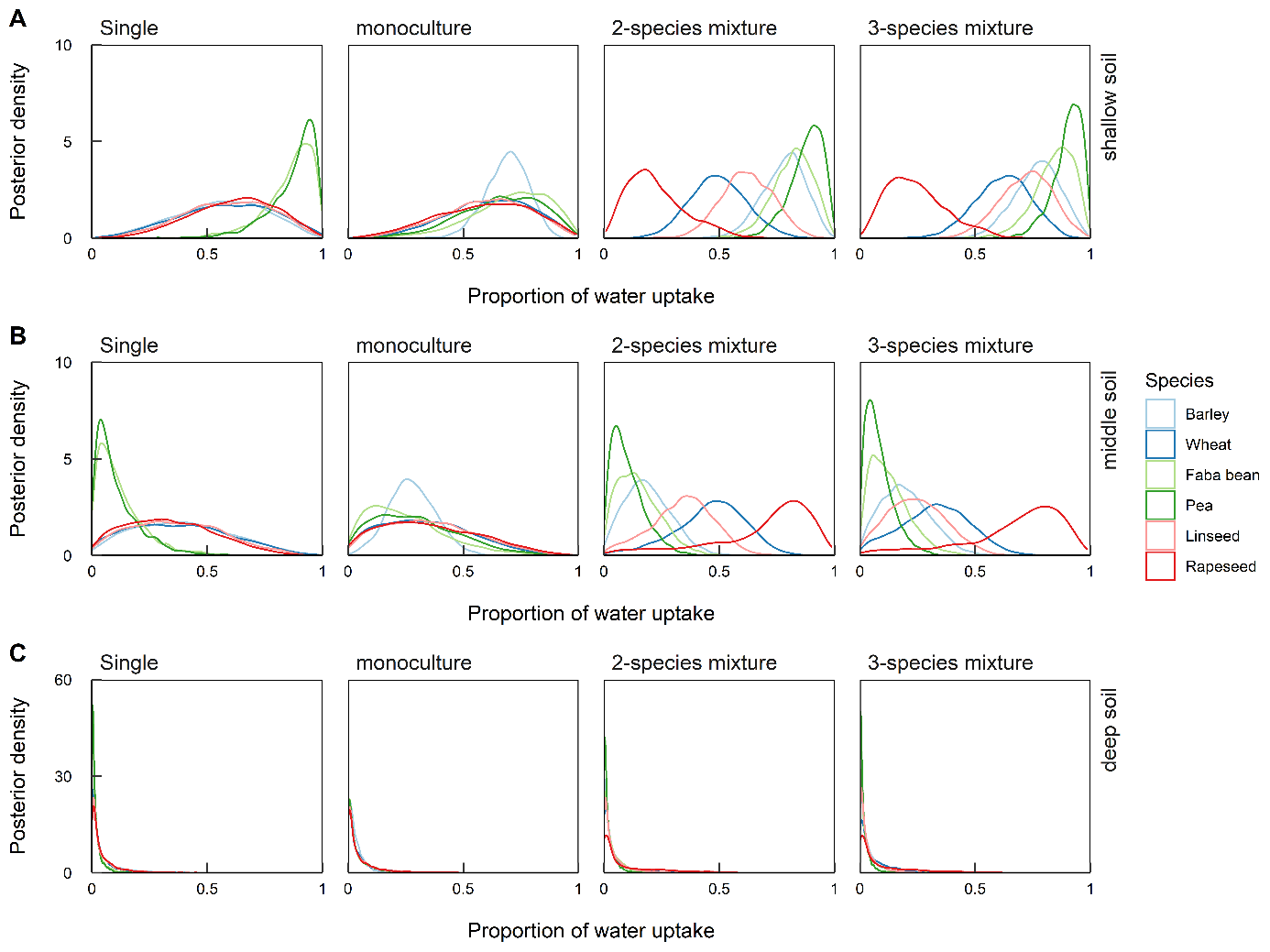

**Fig. S2**. Kernel densities from posterior distributions of water uptake proportions estimated by the mixing model MixSIAR (model 1; species*diversity as fixed factor, species composition as random factor) in the different soil layers (shallow, middle and deep) and in the different diversity levels (single plant, monoculture and 2- and 3-species mixtures) for the six species (coloured lines). Note the different y-axis scales.

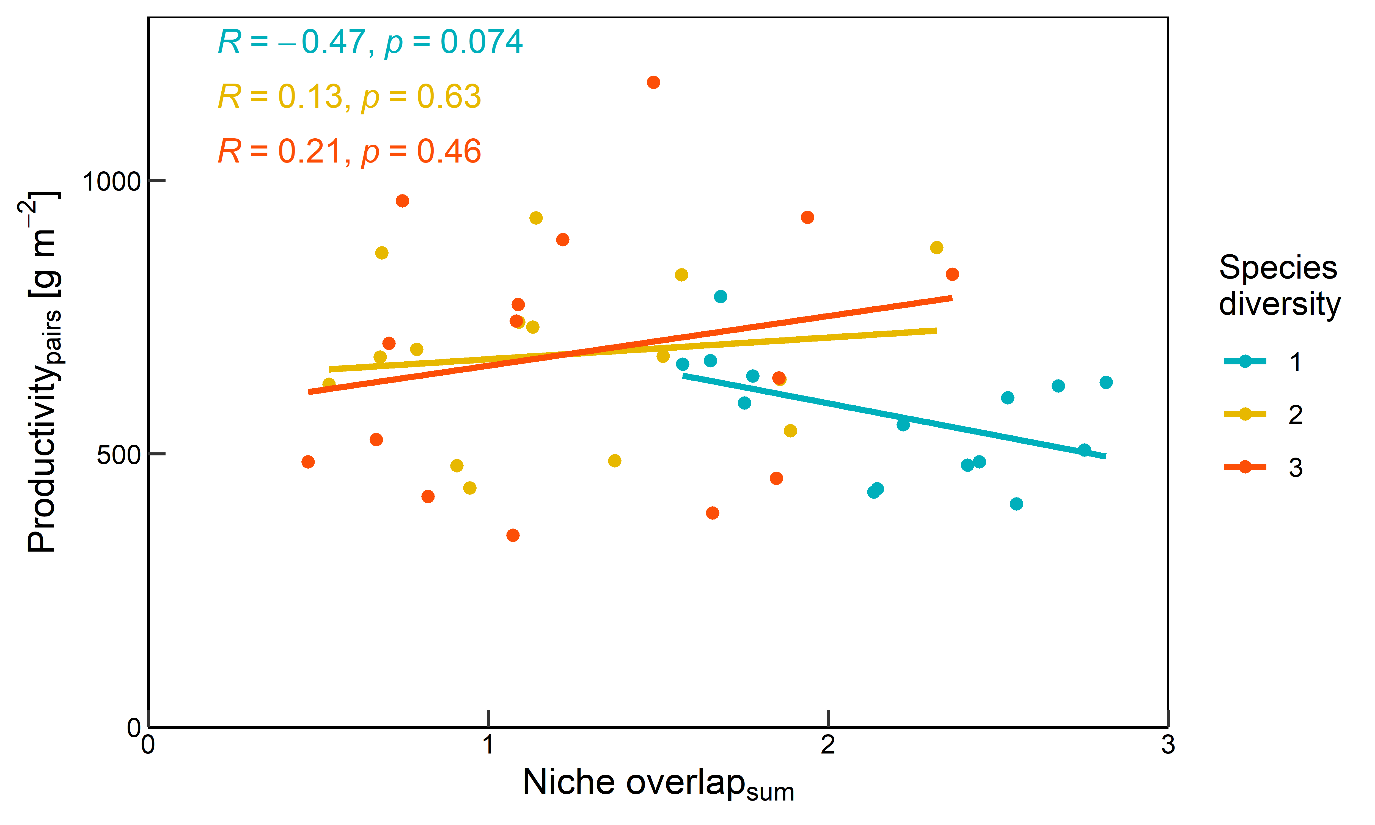

**Fig. S3**. Relationship between productivity and niche overlap of hypothetical species pairs grown in monocultures (blue) and 2- (yellow) and 3-species mixtures (red). The regression line and the results from the Pearson correlation are indicated.

**Table S2**. Type I-analysis of variance of the linear mixed models (LMM) in Fig. 2. The first LMM (relationship between δ^18^O and δ^2^H) the response variable was δ^18^O, the predictor δ^2^H and the random term the plot. In the LMM of δ^18^O and δ^2^H along the soil profile, respectively, the response was the soil depth, the predictors δ^18^O and δ^2^H, respectively, and the random term the plot.

|  |  | **Df** | **denDf** | **F** | **P** |
| --- | --- | --- | --- | --- | --- |
|  |  | **Relationship between δ^18^O and δ^2^H** | | | |
| δ^2^H |  | 1 | 55.8 | 2321 | **<0.001** |
|  |  | **δ^18^O along the soil profile** | | | |
| δ^18^O |  | 1 | 58.1 | 302.20 | **<0.001** |
| δ^18^O^2 |  | 1 | 58.1 | 48.66 | **<0.001** |
|  |  | **δ^2^H along the soil profile** | | | |
| δ^2^H |  | 1 | 59 | 247.60 | **0.011** |
| δ^2^H^2 |  | 1 | 59.3 | 20.48 | **<0.001** |

**Table S3**. Type I-analysis of variance (ANOVA) of the linear mixed model with the square-root transformed relative interaction index (RII) as response variable and block, diversity (monoculture vs mixture) and mixture diversity (2- vs 3-sepcies mixture) as explanatory variable and species composition as random term. Df: degrees of freedom, denDf: denominator degrees of freedom, F: probability distribution, P: error probability. P-values in bold are significant with α=0.05.

|  | **Df** | **denDf** | **F** | **P** |  | **Df** | **denDf** | **F** | **P** |
| --- | --- | --- | --- | --- | --- | --- | --- | --- | --- |
|  | **Barley** | | | |  | **Wheat** | | | |
| intercept | 1 | 7 | 777.60 | **<0.001** |  | 1 | 7 | 640.50 | **<0.001** |
| block | 2 | 18 | 0.96 | 0.400 |  | 2 | 18 | 0.52 | 0.602 |
| diversity | 1 | 7 | 0.01 | 0.917 |  | 1 | 7 | 0.00 | 0.998 |
| mixture diversity | 1 | 7 | 0.27 | 0.618 |  | 1 | 7 | 0.00 | 0.967 |
|  | **Faba bean** | | | |  | **Pea** | | | |
| intercept | 1 | 7 | 2724.00 | **<0.001** |  | 1 | 7 | 1629.00 | **<0.001** |
| block | 2 | 18 | 0.21 | 0.812 |  | 2 | 18 | 0.25 | 0.778 |
| diversity | 1 | 7 | 1.89 | 0.212 |  | 1 | 7 | 0.21 | 0.661 |
| mixture diversity | 1 | 7 | 1.16 | 0.316 |  | 1 | 7 | 3.46 | 0.105 |
|  | **Linseed** | | | |  | **Rapeseed** | | | |
| intercept | 1 | 6.6 | 993.70 | **<0.001** |  | 1 | 7.1 | 88.97 | **<0.001** |
| block | 2 | 17 | 2.13 | 0.145 |  | 2 | 16.1 | 2.38 | 0.122 |
| diversity | 1 | 6.4 | 6.04 | **0.047** |  | 1 | 8.3 | 0.05 | 0.822 |
| mixture diversity | 1 | 6.6 | 6.6 | 0.891 |  | 1 | 7.0 | 0.03 | 0.913 |

**Table S4**. Type III-analysis of variance (ANOVA) of the generalised linear mixed models with the estimated proportions of water uptake (model 4) as response variable (each soil layer separately), the relative interaction index (RII), diversity and their interaction as predictors and species composition as random term. Df: degrees of freedom, Chisq: Chi-square statistic, P: error probability. P-values in bold are significant with α=0.05.

|  | **Df** | **Chisq** | **P** |  | **Df** | **Chisq** | **P** |  | **Df** | **Chisq** | **P** |
| --- | --- | --- | --- | --- | --- | --- | --- | --- | --- | --- | --- |
| **soil layer** | **shallow** | | |  | **middle** | | |  | **deep** | | |
| RII | 1 | 585 | **0.016** |  | 1 | 6.76 | **0.009** |  | 1 | 3.73 | 0.053 |
| diversity | 2 | 1.69 | 0.428 |  | 2 | 1.93 | 0.380 |  | 2 | 1.47 | 0.481 |
| RII × diversity | 2 | 1.27 | 0.531 |  | 2 | 1.34 | 0.511 |  | 2 | 1.26 | 0.533 |
